# A quick and efficient method enabling lncRNA functional studies in the marine diatom *Phaeodactylum tricornutum* using a multiplexed CRISPR-Cas9 editing approach

**DOI:** 10.64898/2026.10.08.756992

**Authors:** Florent Charton, Neo Revillon, Priscillia Pierre-Elies, Hiroaki Matsui, Chris Bowler, Yusuke Matsuda, Helena Cruz de Carvalho

## Abstract

**Background:** Long non protein coding (lnc)RNAs are ubiquitous transcripts known to be major regulators of cellular processes in eukaryotes. In diatoms, specific lncRNA expression is tightly correlated to nutrient fluctuations, which suggests function, but their biological relevance remains to be established. In order to fill this knowledge gap, we sought to design a protocol to generate diatom lncRNA knockout mutants by CRISPR-Cas9 enabling subsequent functional studies to shed light into their mechanistic roles. Since lncRNAs do not contain open reading frames that can be targeted by point mutations for the generation of functional knockouts, we designed a multiplexed CRISPR-Cas9 plasmid harboring two independent gRNA for the specific ablation of lncRNA loci in *P. tricornutum*. As proof of concept, we used our plasmid-based system to target three lncRNAs whose expression is known to be responsive to phosphate depletion.

**Results:** Using the golden gate based uLoop system, we assembled a single plasmid containing all the necessary components to generate CRISPR-Cas9 large genomic ablations of lncRNAs in *P. tricornutum* using bacterial conjugation. Three intergenic lncRNAs (*Linc91*, *Linc77* and *Linc24*) responsive to phosphate fluctuations were selected and independently targeted. Around 32% of the exconjugants presented a monoallelic ablation of the targeted loci and 11% presented a biallelic ablation on the first PCR-screening round. Mutated lines, harboring biallelic lncRNAs ablations for each lncRNA targeted, were successfully generated for each of the targeted lncRNAs. Additionally, a first line of phenotypic analysis revealed significant growth rate differences for one of the mutants (*Δlinc24*) under normal and phosphate depleted conditions, suggesting its involvement in cell division regulation under nutrient fluctuations.

**Conclusions:** This work provides a simple and efficient method for the generation of lncRNA knock-out mutants in *P. tricornutum*. Furthermore, this approach can be easily adapted to other microalgae amenable to transformation, enabling the enhancement of our understanding of these important cellular regulatory transcripts in the marine environment.

## Background

Diatoms (*Bacillariophyceae*) are one of the dominant eukaryotic phytoplankton groups in the modern ocean. Despite their microscopic size and representing less than 1% of the photosynthetic biomass on Earth, they are responsible for ∼45% of the ocean primary productivity [1]. Diatoms are known to be excellent competitors, especially in turbulent areas subjected to nutrient fluctuations. Under these environments where nutrients are supplied with high pulse frequencies, diatoms are known to outcompete other phytoplankton groups by depriving them of these essential resources [2]. They typically divide quickly, forming large blooms in a few days [3]. When nutrients are lacking, diatoms rapidly adjust their metabolism to halt growth until favorable conditions are again met, when they efficiently resume cell division [3–5]. This capacity is ultimately underpinned by a sensory and regulatory system that is fine-tuned to detect environmental changes and regulate molecular pathways, allowing diatoms to quickly and efficiently optimize their metabolism to the ocean’s changing environment. However, the molecular underpinnings of this response, providing diatoms with an extreme resilience to environmental fluctuations, remains elusive.

For a long time considered as artifactual transcriptional noise, long (>200 nt) noncoding RNAs (lncRNAs) are now recognized to be major regulators of cellular processes in eukaryotes [6–8]. LncRNAs, like messenger RNAs (mRNAs), are mostly transcribed by RNA polymerase II, being capped, polyadenylated and even spliced [6, 9]. However, unlike mRNAs, they are active as RNA molecules and generally do not produce functional polypeptides. Regardless of being named for what they do not do [6], lncRNAs have been shown to be involved in almost every step of gene regulation, having key roles during cellular differentiation, development, and in response to stress [6, 10–13]. Due to their intrinsic molecular nature, lncRNAs are known to interact with a multitude of other molecules, from DNA to proteins to other RNA transcripts, both in the nucleus and the cytoplasm. In the nucleus, lncRNAs have been reported to interact with histone-modifying complexes regulating changes to the chromatin landscape, to interact with transcription factors and/or RNA polymerase II, and to regulate RNA maturation and stability [6, 13]. In the cytoplasm, lncRNA have been reported to be involved in regulating miRNA activity, by acting as molecular sponges and titrating their concentration, a process known as target mimicry [13, 14] and to regulate enzymatic activity, namely in post-transcriptional modifications like phosphorylation, ubiquitination and SUMOylation [15].

Although the central regulatory roles of lncRNAs have been extensively described essentially in mammals, hundreds of thousands of lncRNAs have been identified across the Eukarya domain of life, from animals to plants, to yeasts and unicellular protists [4, 16–20]. In the marine diatom *Phaeodactylum tricornutum*, over 40% of the processed transcripts under phosphate fluctuations lack protein-coding capacity [4, 21]. These are polyadenylated *bona fide* transcripts longer than 200 nucleotides which can be classified as lncRNAs. *P. tricornutum* lncRNAs have lower GC content and lower free energy compared to mRNAs [4, 21], traits which are commonly found in functional human lncRNAs, which can be related to their lower expression levels and high interaction capacity [9, 22]. This reveals cross-kingdom feature conservation which supports functionality for diatom lncRNAs. Furthermore, *P. tricornutum* lncRNAs are not expressed randomly but rather under specific environmental conditions, namely under phosphate stress [4, 21], high *p*CO2 [23], hyposalinity [19] and high light [24], further supporting function. However, the biological relevance of diatom lncRNAs remain to be demonstrated.

A key approach to investigate the role of genes of unknown function in a biological context is to specifically knockout the transcript and study the resulting phenotype. At present, one of the most efficient and simple ways to achieve this is through the Clustered, Regularly Interspaced, Short Palindromic Repeats, (CRISPR) associated with the Cas9 nuclease, which allows the precise targeting and editing of a specific genomic locus. The CRISPR-Cas9 methodology has been widely adopted to knockout (KO) protein coding genes, where a point deletion or insertion will usually result in a frameshift in the open reading frame (ORF) leading to a precocious STOP codon, which changes the polypeptide sequence and length [25]. The CRISPR-Cas9 has already been successfully used in *P. tricornutum* to stably modify its genome [26–32]. However, this approach has never been used to target non-protein coding genes in diatoms, namely lncRNAs and, to our knowledge, neither in any other microalgae. The undertaking of functional studies using the CRISPR-Cas9 approach by the generation of KO mutants is more challenging for lncRNAs than for mRNAs because, unlike the latter, lncRNAs lack the open reading frames (ORFs) that are usually targeted. Since a simple point mutation will most likely not lead to a loss of function, a powerful approach to uncover lncRNA function would be to ablate its genomic locus [33].

With this in mind, we designed a multiplexed CRISPR-Cas9 plasmid harboring two independent guide (g)RNAs with the aim to ablate the lncRNA genomic locus to target in order to generate *ΔlncRNA* diatom mutants. As proof-of-concept, we chose to target three *P. tricornutum* lncRNAs previously shown to be responsive to phosphate fluctuation [4] to target. To avoid multiple copy integration and excess Cas9 in the cells that could lead to increased probability of off-target activity, we opted to deliver the recombinant plasmid as an episome by bacterial conjugation, which can be curated once the mutation is confirmed and stable. Our results show that our method successfully generated diatom mutants harboring biallelic ablations of the targeted lncRNAs. This represents an important starting point that will enable subsequent functional studies aimed at shedding light into the roles and mechanistic function of lncRNAs in diatoms.

## Materials and Methods

### Culture conditions

Axenic cultures of *P. tricornutum* strain Pt1 (CCMP632) obtained from the Center for the Culture of Marine Phytoplankton (Maine, USA) and derived mutants were cultured in 50 ml f/2 media [34] filtered steam-sterilized artificial seawater (40 g/L, NutriSelect® Basic, Merck Millipore) supplemented with f/2 nutrients, elements and vitamins (f/2 Media Kit, Bigelow) with the exception of silica (f/2-Si), in tissue culture flasks (75 cm^3^) with vented filter caps (CytoOne, Starlab) under continuous shaking (90-100 rpm) at 19°C, under cool white fluorescent lights at ∼60 µmol.m^-2^.s^-1^, with a 12 h photoperiod.

### Choice of lncRNA loci to target and design of single guide RNAs

Specific single guide (g)RNAs for the Cas9 recognition sites in the targeted lncRNA loci were designed using both the CRISPOR [35] and the PhytoCRISP-Ex [36] software. Two gRNAs were designed for each lncRNA locus, one targeting its 5’end and one targeting its 3’end.

### Plasmid construction

All plasmids used in this work, with the exception of the L1_2_hCas9 plasmid, were generated using the recursive Golden Gate-based uLoop system [37]. The final conjugation Level 2 plasmids used to transform *P. tricornutum* by bacterial conjugation were assembled using four Level 1 plasmids and the pCae1 receiver plasmid [37]. The four plasmids included the L1_2_hCas9 plasmid, generated by the amplification of the Cas9 gene from the plasmid pDEST-HA-hCas9 [38]. Adapters to perform the NEBuilder HiFi DNA assembly were added to the sequence, allowing its integration in the pCao2 backbone [37]. NEBuilder HiFi DNA assembly was performed following the manufacturer’s instructions and final assembly was verified by Sanger sequencing. Also included was the plasmid pSBL1-1-PTCv2 which contained all the necessary elements for the plasmid maintenance in *P. tricornutum* [39]. Two plasmids, L1-3_gRNA and L1-4_gRNA, despite being almost identical use the uLoop L1-3 and L1-4 adaptor sequences backbones, respectively, and were used for the two different gRNA of each final L2 plasmid. The initial gRNA sequences were amplified from the pU6::AOX-gRNA [38] with adapters to perform a NEBuilder HiFi DNA Assembly and being integrated in pCao3 et pCao4 respectively. NEBuilder HiFi DNA was performed following the manufacturer’s instructions and the final assembly was verified by sequencing.

All plasmids were amplified by transformation in One Shot® Top10 Chemically Competent *E. coli* cells by heat shock (as described above) and purified by a Miniprep using the kit Macherey-Nagel® NucleoSpin plasmid and following the manufacturer’s instructions. Purified plasmids were resuspended in nuclease-free water (Qiagen) and profiled using restriction enzymes to ensure plasmid integrity.

### gRNA plasmid substitution

Designed gRNAs (two per lncRNA locus to be targeted) were inserted in the final plasmids (L1-3 and L1-4) by PCR. Briefly, a PCR was made using primers containing the target gRNA sequence, followed by GTTTTAGAGCTAGAAATAGC (forward primer) and the reverse gRNA sequence followed by CGACTTTGAAGGTGTTTTTTG (reverse primer), which were specific to the plasmid site where the new gRNA was to be integrated. The PCR product was then digested by the DPN1 enzyme to cleave the original template plasmid and directly transformed into One Shot® Top10 Chemically Competent *E. coli* as described above. All plasmids were sequenced following a Miniprep purification, as described above.

### Transformation of *P. tricornutum* by bacterial conjugation

The transformation protocol used was an adaptation of the protocol published by [39]. Four days before the conjugation, 5.10^6^ *P. tricornutum* cells from cultures grown in f/2-Si media in exponential phase were plated in 45% ESAW 5% LB agar in 12 well-plates and kept in the growth culture chamber for 4 days. Three days before the conjugation, *E. coli* cells containing the pTA-MOB helper plasmid, needed for the transfer of DNA construct form the bacteria to a recipient cell, were transformed by heat shock (same protocol as above) with our plasmid of interest and then selected on plate for both plasmids using gentamicin (20 mg.l^-1^) for the pTA-MOB plasmid selection and spectinomycin (30 mg.l^-1^) for the L2 level final multiplexed CRISPR/Cas9 assembly plasmid. The day before the conjugation, transformed bacteria were pre-cultured in liquid LB selective media at 37° C overnight. On the morning of conjugation, pre-cultured bacteria were refreshed in LB selective media and incubated at 37° C for 2-3h (to reach exponential growth phase) until DO600 = 0,6-0,8. Bacteria were then pelleted and gently resuspended in 50 µl LB (without antibiotics) for every 5 ml of harvested culture. A 50 µl bacteria culture was added in each well of 4-day old plated *P. tricornutum* cells. The plates were then kept in the dark for 1h30 at 30° C and then moved for 3 days to the growth chamber (under photoperiodic light). After 3 days, *P. tricornutum* cells were resuspended in 500 µl ESAW liquid media and 250 µl was plated in fresh 50% ESAW - 50% Agar selective plates containing phleomycin (transformant *P. tricornutum* selection) and Ampicillin100, Streptomycin100 and Chloramphenicol34 in order to kill bacteria from the conjugation and avoid further contamination (two plates were made for each conjugation).

### Isolation of bi-allelic *ΔlncRNA* knock-out mutant lines

Two weeks after conjugation, up to 96 exconjugant colonies were transferred from the conjugation plates to fresh selective plates. After 1 week growth, cells were resuspended in 30 µl ESAW or f/2 media, from which 15 µl was plated in a replicated selective plate and the other 15 µl was used to perform DNA extraction for PCR screening to verify the occurrence of the CRISPR-Cas9 ablation of the targeted locus. At least three independent positive colonies from the PCR screening for the CRISPR-Cas9 lncRNA locus ablation were selected per conjugation plate (biological replicates). These selected positive colonies were resuspended in 40 µl ESAW (or f/2) media, from which 15 µl was used to perform DNA extraction and a new PCR screening, 15 µl were replicated in a new selective plate (back-up) and the remainder 10 µl used for dilutions in a final volume of 500 - 1000 µl. Following the dilutions, volumes between 100 - 300 µl were plated in a fresh selective plate. After 2 - 3 weeks growth, up to 5 colonies per biological replicate (minimum 15 colonies per lncRNA) were picked and resuspended in 40 µl ESAW (or f/2) to be plated in a replicated selective plate and to perform a screening PCR as in the previous step. Ten µl were used to start a liquid culture with a final volume of 100 µl selective f/2 (or ESAW media) in 12 well-plates. At this step, the PCR provided information on the purity of the colony (putatively chimeric or heterozygous/monoallelic for the full lncRNA ablations) selected on the previous steps, and enabled the identification of isolated clones with bi-allelic mutations for the CRISPR-Cas9 full ablation. The liquid preculture (100 µl) of the isolated mutant lines were then used to progressively increase the volume of the cultures. In order to eliminate the episomes from the confirmed bi-allelic mutant lines, the pressure selection was removed after one or two cycles of culture refreshing.

### DNA extraction for colony PCR

DNA extraction was performed using 10 µl of resuspended cells from a plate or a pellet of 1ml liquid culture after centrifugation at 10000 *g* for 1 min. Lysis was performed by adding 10 µl of NP-40 lysis buffer (NaCl 150mM, Tris-Cl 50mM pH 8, NP-40 1%) for 10 mins at 95°C. After a short centrifugation (10000 *g*, 1min), 10 µl of the supernatant was diluted in 90 µl DNAse/RNAse free H_2_O and this DNA extract was used for the following PCR.

### PCR genotyping of the mutant lines

Genotyping of exconjugants was performed using the DreamTaq Green PCR Master Mix and specific primers of the region of interest (Supplementary Table S1). The default PCR program was the following: 55°C for the annealing T° (changed/optimized if no PCR amplification was obtained) and the elongation time was adjusted to 45 sec/kb. PCR products were separated by agarose gel (1-1.5%) electrophoresis using ethidium bromide and analyzed with UV after migration in Tris-acetate-EDTA (TAE) buffer (Ogden and Adams, 1987). For sequencing purposes, the PCR amplification was performed using Phusion Green Hot start II, using the same program as described above. PCR products were loaded on an agarose gel as described above and the separated DNA bands were cut out from the gel and purified using the Nucleospin Gel and PCR Clean-up (Macherey-Nagel) following the manufacturer’s instruction. Sanger sequencing was performed by Eurofins Genomics.

### Growth responses to inorganic phosphate fluctuations

Inorganic phosphate (Pi) fluctuation experiments were performed as described in [4]. Briefly, equal aliquots of 4-d-old cultures were inoculated in parallel in f/2-Si media with (control) or without phosphate supplement (Pi depleted) under the same growth conditions as above. Pi resupplementation was made on 4-d-old Pi depleted cultures by pelleting the cells at 1500 *g* for 10 min at 4° C and resuspended in the same volume of fresh full f/2-Si media. Culture growth was followed daily or every two days, using a hemocytometer (Fisher Scientific, USA) or with a Luna automated cell counter (LUNA-FX7™). Specific growth rates (μ, day^-1^) were calculated during exponential growth using the following formula: μ = ln(N1/N0)/(t1-t0) where N1 and N0 are the cell densities at the end (t1) and of the beginning (t0) of the exponential growth phase, respectively.

## Results

### *P. tricornutum* lncRNAs candidates selected for CRISPR-Cas9 editing

The lncRNA candidates to target by the multiplexed CRISPR-Cas9 approach were selected according to the following criteria: 1) to be significantly differentially expressed under phosphate (Pi) fluctuations [4], 2) the lncRNA had to be intergenic (i.e., its genomic locus should not overlap with any protein coding gene, both in sense and antisense) to limit potential side effects of a simultaneous deletion of a protein coding gene on the opposite strand by the Cas9 endonuclease and, 3) be a strong lncRNA candidate based on a VotingLNC lncRNA probability score of 0.6 or higher [20]. The candidate lncRNAs that followed these criteria were then used to undertake a BLAST analysis on Phatr3 genome annotations to make sure that the transcribed locus of each lncRNA did not overlap with a newly annotated protein coding gene or a putative regulatory sequence (i.e., a putative promoter region) and were present as a single copy in the genome. A BLAST analysis was also performed on the long-read contig sequences from [40] to further confirm the absence of multiple copies. After this screening, we selected three lncRNAs to perform the proof-of-concept multiplexed CRISPR-Cas9 ablation, named *Linc24*, *Linc77* and *Linc91* (Table 1). *Linc24* is a 502 nt long intergenic lncRNA located in chromosome 10 (chr_10:232961-233462 [-]). It has a VotingLNC lncRNA potential score of 0.8 and is upregulated under Pi depletion. *Linc77* is a 454 nt long intergenic lncRNA located in chromosome 5 (chr_5:742240-742694 [+]). It has a VotingLNC lncRNA potential score of 1 and is downregulated under Pi depletion. *Linc91* is a 906 nt long intergenic lncRNA located in chromosome 23 (locus chr_23:364524-365429[-]). It has a VotingLNC lncRNA potential score of 0.8 and, like *Linc24*, it is upregulated under Pi depletion. All three lncRNAs are therefore strong lncRNA candidates for functional studies, using the CRISPR-Cas9 approach.

**Table 1.** *P. tricornutum* lncRNA candidates selected for CRISPR-Cas9 genomic locus ablation.

| Name | Locus | Length (nt) | DE.P4d | DE.P8d | DE.R4d | VotingLNC score |
| --- | --- | --- | --- | --- | --- | --- |
| *Linc24* | chr_10:232961-233462 [-] | 502 | U | U | D | 0.8 |
| *Linc77* | chr_5:742240-742694 [+] | 454 | D | – | D | 1 |
| *Linc91* | chr_23:364524-365429 [-] | 906 | U | U | – | 0.8 |

DE, differential expressed; P4d, phosphate stress 4 days; P8d, phosphate stress 8 days; R4d, phosphate stress recovery 4 days; U, upregulated; D, downregulated (DE cutoffs were defined by a two-fold variance in FPKM and a *P* value (Fisher's exact test) of < 0.05, data collected from [4]).

### LncRNA genomic ablation strategy

In this work our goal was to generate diatom mutants harboring a complete knockout of the selected lncRNAs. To do so, for each lncRNA we designed two specific gRNAs flanking the 5’ and 3’ ends of the corresponding loci, which would lead to the ablation of the region between the position of both guides by the Cas9 enzyme. The gRNAs were designed using PhytoCRISP-Ex [36] and CRISPOR [35]. For each lncRNA, we selected pair of gRNAs presenting no mismatches in the seed sequence and which allowed for the CRIPSR-Cas9 ablation of at least 50% of the targeted loci in order to generate true functional knockouts. In the case of *Linc24*, the selected gRNAs targeted a region corresponding to approximately 87% of its assigned genomic locus, for *Linc77*, it corresponded to ∼60% and for *Linc91* to ∼90% (Fig. 1).

**Fig. 1.**
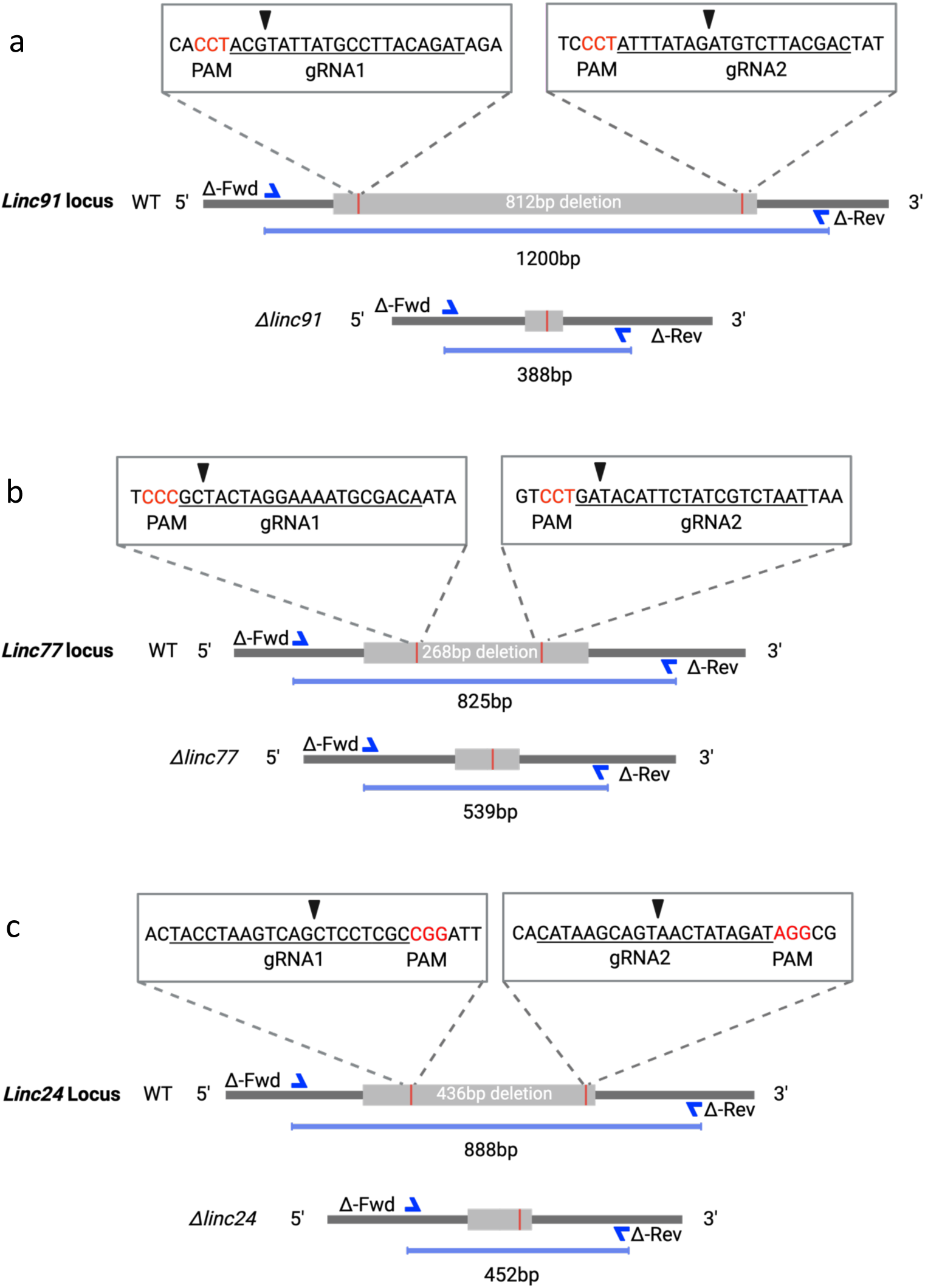
Genomic ablation of lncRNA loci by CRISPR-Cas9 in *P. tricornutum*. **a-c** Schematic representation of the position of each gRNA couple on the loci of the three targeted lncRNAs (**a** *Linc91*; **b** *Linc77*; **c** *Linc24*). The 20-nt region targeted is underlined, the PAM (Protospacer Adjacent Motif) sequence is in red and the arrows points to the Cas9 nuclease cutting position. The position of the primer pairs (forward, ΔFwd and reverse ΔRev primers; Supplemental Table S1) used to screen the exconjugant DNA by PCR then sequencing with the expected amplification size (in bp) in both the wild-type (WT) and the (Δ) mutants is also shown. Diagram created in https://BioRender.com.

### CRISPR-Cas9 multiplexed plasmid design

CRISPR-Cas9 multiplexed plasmids were constructed modularly using the uLoop Golden Gate-based assembly system, which allows the assembly of multiple fragments without the use of a PCR or fragment isolation [37]. Four Level 1 (L1) plasmids were initially prepared, each containing the needed “domesticated” sequences (Fig. 2). The advantage of the uLoop system is its recursiveness based on the “even and odd” levels and the four plasmids used were in either an “even” or “odd” vectors. We domesticated the hCas9 with the *FcpB* promoter into an L1 plasmid (pL1_2_hCas9) as well as the two gRNAs for each lncRNA, each directed with the pU6 promoter (L1_3_gRNA/L1_4_gRNA). These were assembled with the core L1 plasmid containing a bacterial origin of replication (pBR322 ORI), a selection marker for bacteria with its promoter (*KanR*), a selection marker in *P. tricornutum* (Pt *shBle*), a eukaryotic origin of replication (oriT), the centromere-like sequence from yeast (CEN_ARS_HIS), which enables the autonomous replication and episome maintenance in the diatom (Karas et al., 2015; Diner et al., 2016), and the receiver pCA_even-1-backbone plasmid (Fig. 2). The resulting final conjugation Level 2 (L2) plasmid, L2-1-Cas9system-Conjugation (12,886 bp) had all the necessary elements to perform transformation by bacterial conjugation in *P. tricornutum*, leading to a CRISPR-Cas9 large ablation of the (lncRNA) targeted locus. Each lncRNA had a L2 plasmid tailor-designed with a locus-specific pair of gRNAs. We also designed a L2 plasmid containing every L1 part described above, minus the L1_3_gRNA/L1_4_gRNA parts (the Cas9 control).

**Fig. 2.**
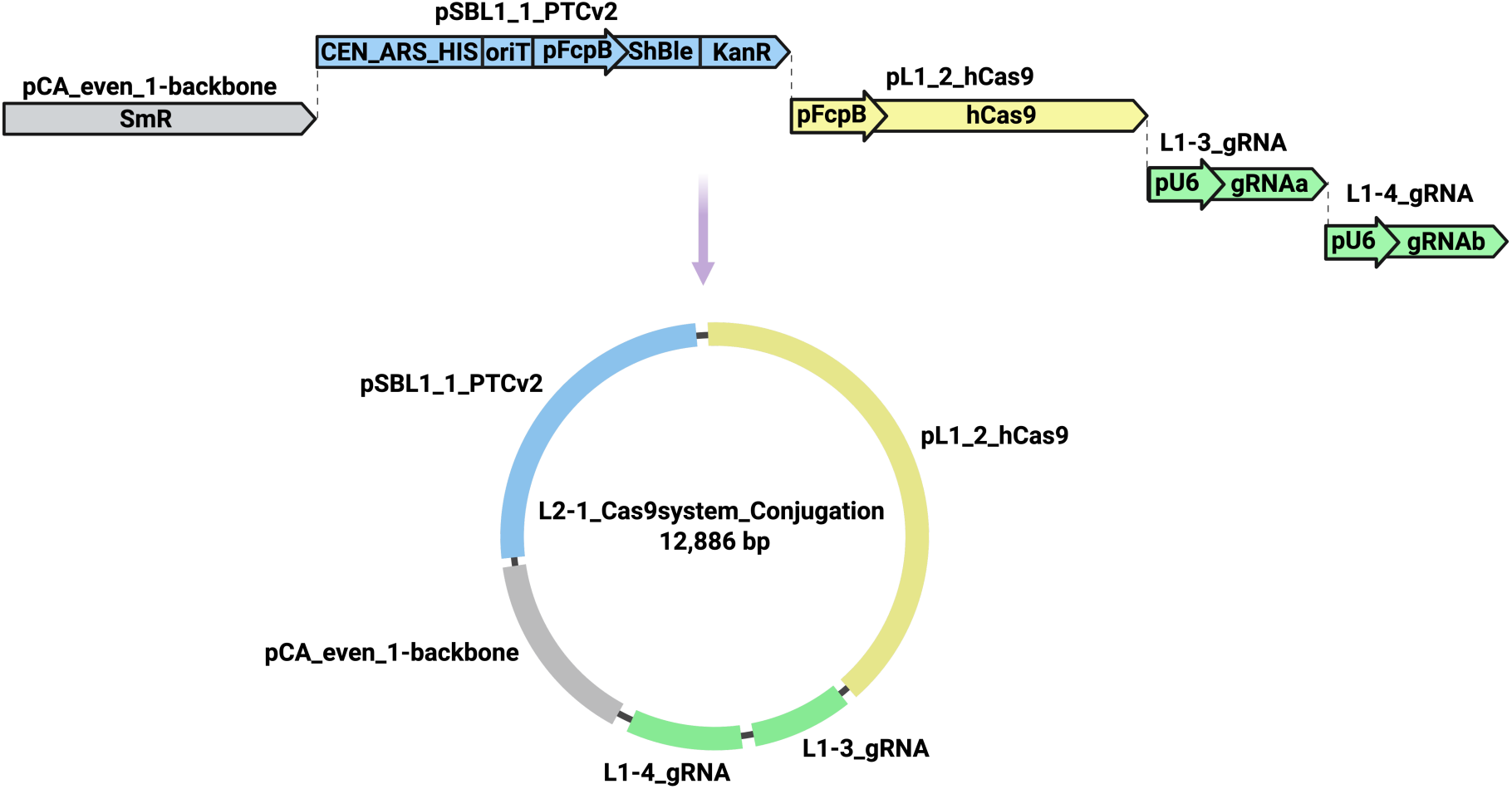
Schematic diagram of the assembly of the multiplexed clustered regularly interspaced short palindromic repeats-CRISPR-associated protein 9 (CRISPR-Cas9) plasmid for lncRNA locus ablation in *P. tricornutum* by bacterial conjugation. The four different Level 1 plasmids were assembled using the uLoop Golden Gate system. Final L2 plasmid (12,886 bp) containing the CRISPR-Cas9 (pFcpB::Cas9), two gRNAs (pU6::gRNAa/b), the selection marker and bacterial origin of replication (pCA_even_1 – Backbone) and the episome maintenance sequences (CEN-ARS-HIS). Diagram created in https://BioRender.com.

### Identification of the CRISPR-Cas9 lncRNA loci ablation events

Following transformation of *P. tricornutum* by bacterial conjugation, exconjugant cell colonies started to be visible after 1-2 weeks. Before transferring them to liquid culture, a colony PCR was performed in order to identify the lines with a complete deletion of the targeted locus. PCR screening of exconjugants using the primer pairs designed for each of the three lncRNA loci (Fig. 1 a-c; Supplementary Table S1), allowed us to identify colonies presenting different editing events (Fig. 3). We found a large variability in terms of putative mutations obtained, namely the absence of detectable ablation, putative monoallelic (either heterozygous or chimeric mixed colonies with WT) and biallelic mutant colonies. Since our goal was to isolate colonies with the full ablation of the lncRNA targeted region, we dismissed any colony that did not reveal the predicted deletion by PCR screening (i.e., ablation not detected), without characterizing further the occurrence and nature of the editing events that might have occurred on these lines. This corresponded approximately to more than half (58%) of the exconjugant cell colonies that appeared on the plates (Fig 3 a). Around 31% of the screened exconjugant colonies (19 out of 62) presented a monoallelic ablation and 11% of the colonies revealed a biallelic mutation (7 out of 62) on the first PCR screening (Fig. 3 a; Supplementary Figure S1). Regarding the putative heterozygous colonies, differences on PCR band intensities suggested chimeric colonies (i.e., WT and mutant) (Supplementary Figure S1). To check this, we diluted the colonies, plated them and, once isolated colonies appeared, performed a new colony PCR genotyping (Fig. 3 b). This was an important step needed to verify that the initial colonies diluted from the same initial clone originated from a single cell and, if not, successive dilutions enabled us to isolate and characterize the different genotypes.

**Fig. 3.**
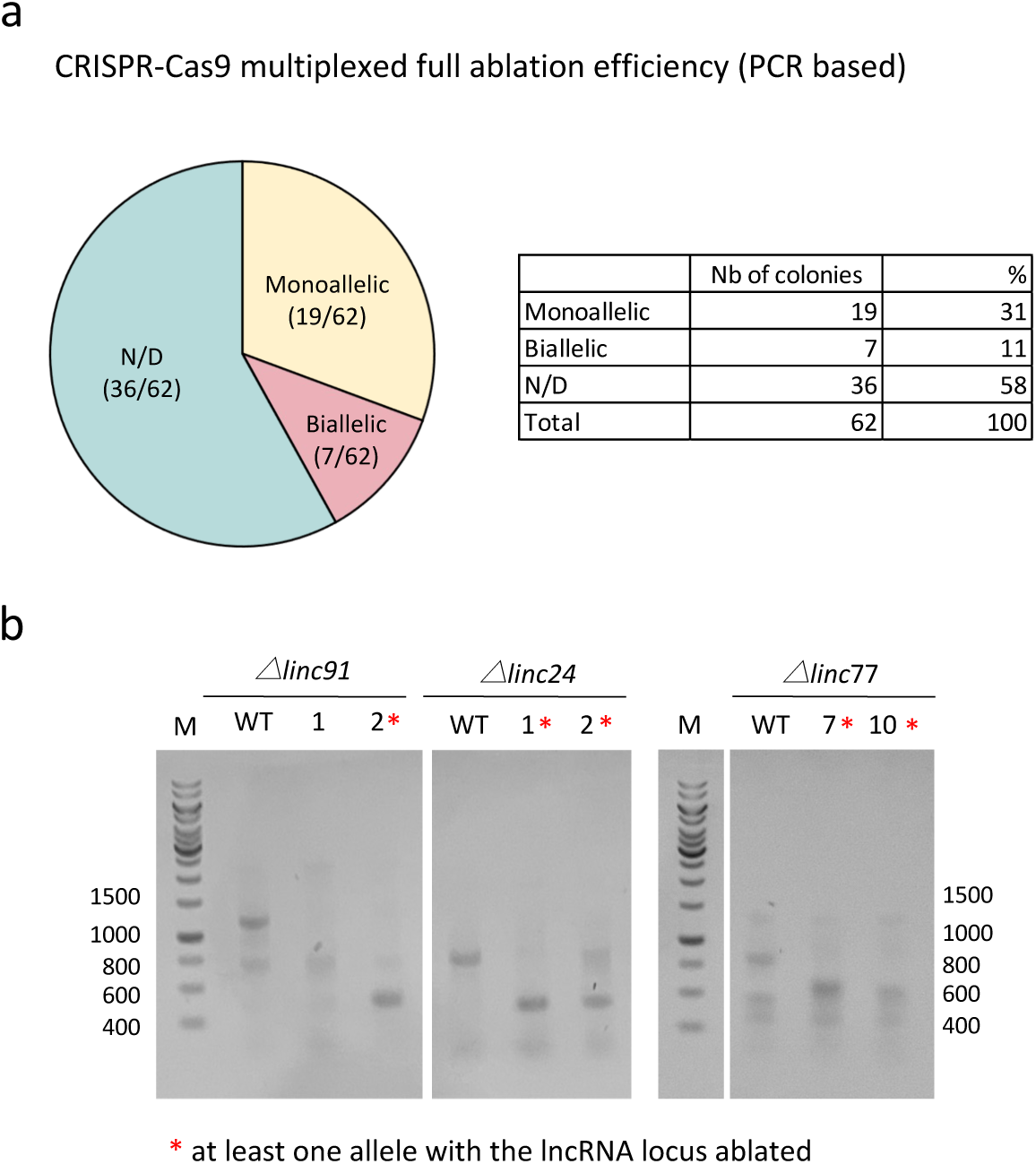
Ablation events in *P. tricornutum* exconjugants analyzed by PCR. **a** Example of the frequency of the multiplexed CRISPR-Cas9 full ablation of the targeted lncRNA loci on both alleles (Biallelic, pink), on one allele (Monoallelic, yellow) and not detected (ND, blue). **b** PCR screening of DNA from exconjugant colonies after the first round of screening and colony isolation/dilution revealing mutant lines presenting at least one of the alleles with the full lncRNA locus ablation (confirmed by Sanger sequencing, Supplementary Figure S2).

### Isolation of biallelic *ΔlncRNA* mutant lines

Following the mutagenesis event and obtaining exconjugant colonies on plates, the first step of the biallelic mutant isolation protocol was to select several colonies from the original plate and replate them in order to obtain more biomass (at each step a replicate plate was made for subsequent analysis). A colony PCR was made on each clone in order to obtain a first information about the genotype (1^st^ verification). In the colonies presenting a mutant band (with the expected deletion) a second PCR was performed to confirm the genotype. These were then submitted to serial dilutions in liquid media and then plated. This liquid dilution allowed to obtain a lower number of colonies, well separated on the plate. At this step we assumed that most of the colonies were derived from only one (or two) cell(s), hence increasing the rate of genetic dependency, meaning that most of the colonies originated from the same initial mutagenesis event (i.e., not actual biological replicates). The final step of the protocol was to genotype by PCR these diluted colonies on plates in order to enable the identification of at least one colony per plate with a single clean mutant band, supporting the presence of a biallelic mutation. From each diluted plate we selected one to two clones for each target lncRNA, and their genomically independence was verified by Sanger sequencing (Supplementary File 1).

### Growth assessment of the mutant lines under nutrient fluctuations

A comparative analysis of the growth responses between the *P. tricornutum* CRISPR-Cas9 KO mutants, the Cas9 control line and the WT lines, was performed under Pi fluctuations in order to have a first line of functional characterization of the mutated lncRNA loci. For all the conditions tested (Full media, Pi depleted and Pi stress recovery), no significant growth differences were found between the WT and the Cas9 line (Fig. 4). Although under full conditions the *Δlinc91* and *Δlinc77* mutant lines seemed to grow slower than the WT, namely around day 6 (Fig. 4 a), no significant differences were detected in their specific growth rates during the exponential phase (between days 2 to 6) (Fig. 4 d). However, significant differences were detected in the specific growth rate of the mutant line *Δlinc24,* which divided significantly faster than the WT line (*P* < 0,01) under normal growth conditions (Fig. 4 d). Under Pi depleted conditions, neither *Δlinc91* nor *Δlinc77* mutant lines show significant growth rate differences from WT (Fig. 4 b, e). However, the *Δlinc24* mutant line presented a significantly higher growth rate than the WT (*P* < 0,05), which was the same trend of what was seen under full growth media. When Pi was re-supplied to the media after 4 days of Pi depletion, none of the mutant lines revealed significant growth rate differences (Fig. 4 c, f).

**Fig. 4.**
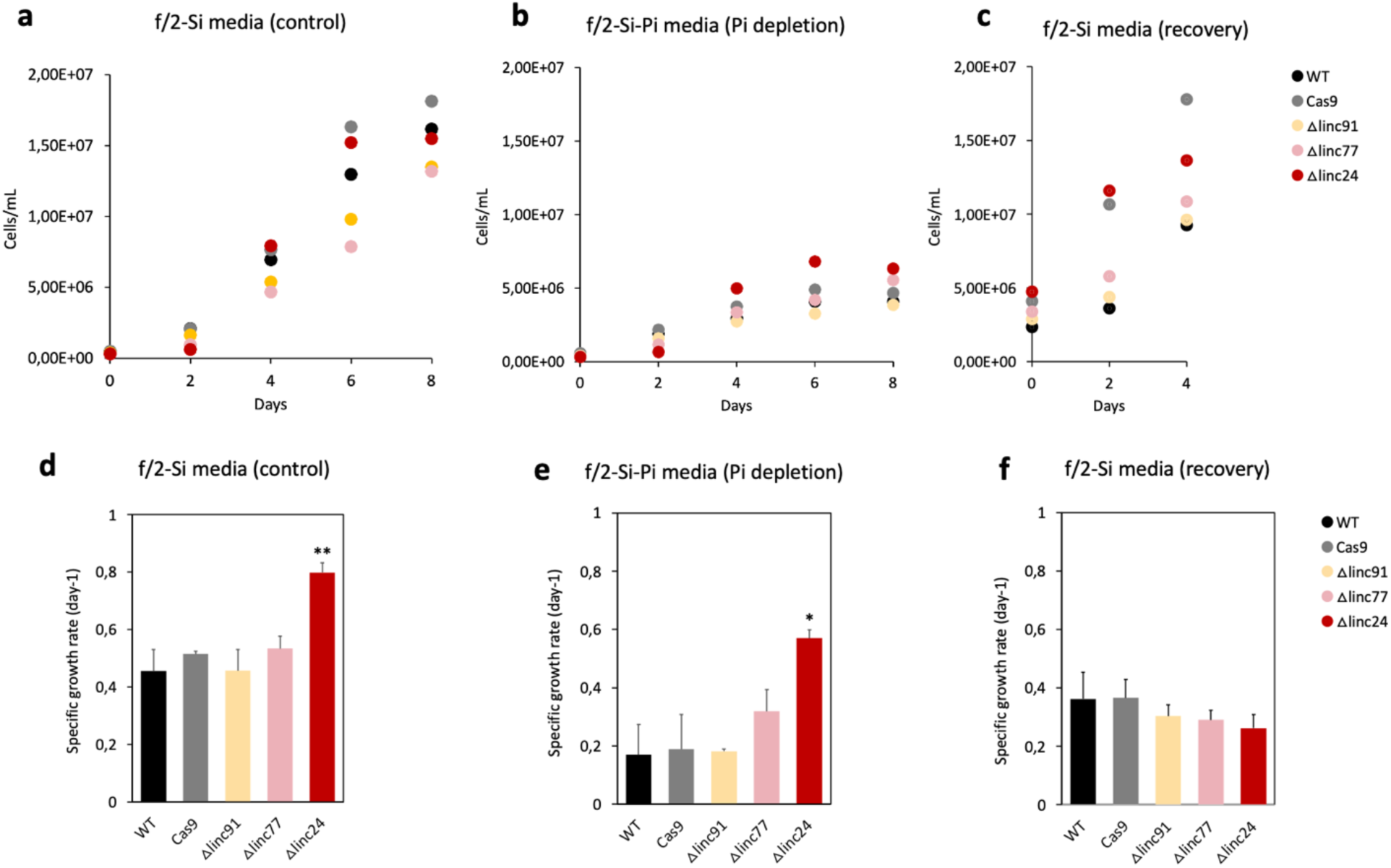
Comparative growth experiments under phosphate fluctuations of the multiplexed CRISPR-Cas9 lncRNA ablated mutant lines, the Cas9 control and the WT lines. **a-c** Culture growth experiment example, and **d-e** Specific growth rates during the exponential phase, of the lncRNA biallelic ablated mutants (*Δlinc91*, *Δlinc77*, *Δlinc24*), the WT and the Cas9 lines. Experiments were made over the course of 8 days in full f/2 media without silica (**a, d**), under inorganic phosphate depleted media (**b, e**) and under four days recovery after four days Pi-depleted (**c, f**). Statistical analysis was performed using a paired-sample Student’s *t*-test with WT as reference: *, *P*<0,05; **, *P*<0,01. Data are the average of at least three replicate measurements for each line, with error bars showing standard deviation.

**Fig. 5.**
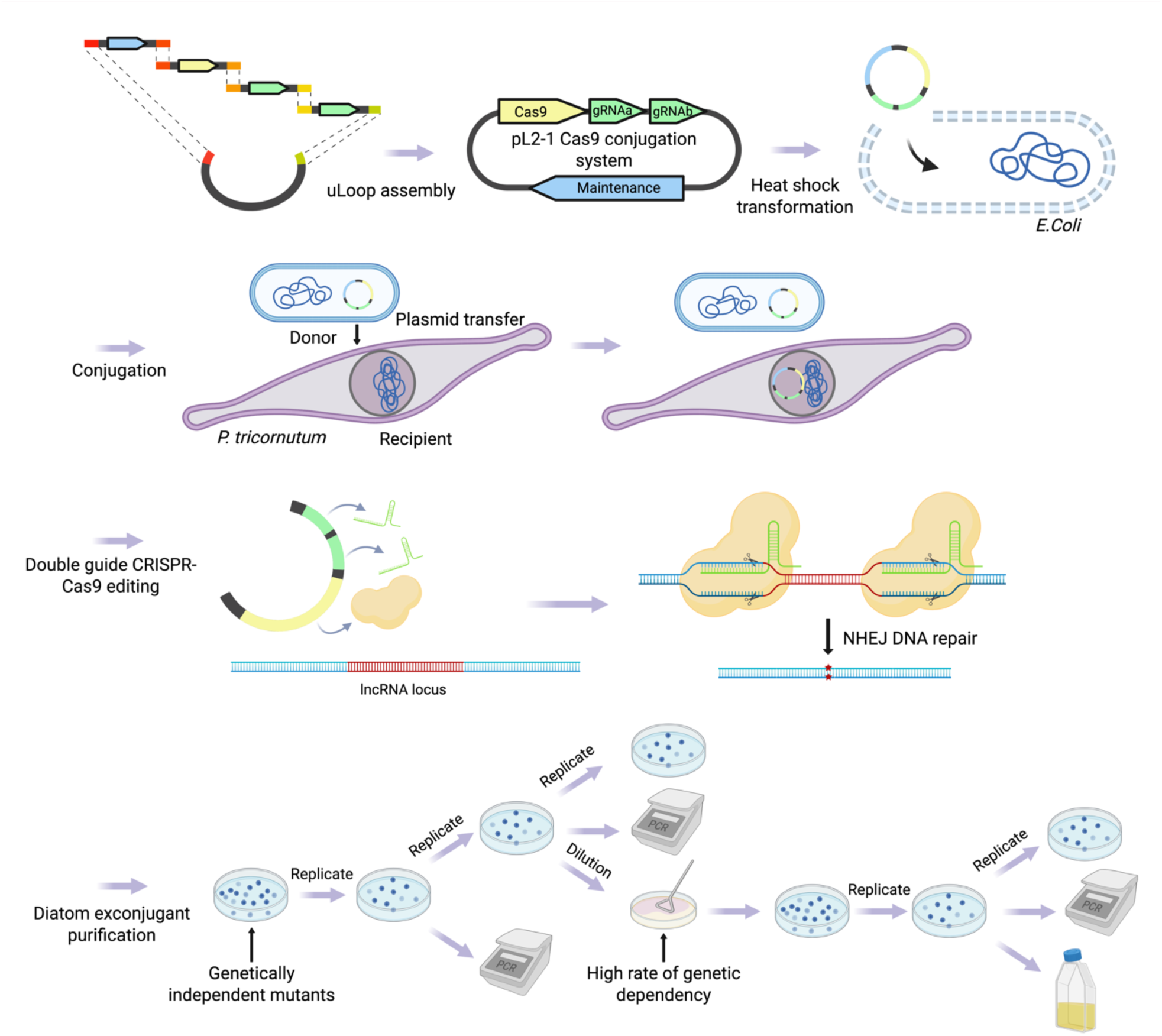
Experimental workflow for CRISPR-Cas9 genomic ablation of lncRNAs in *P. tricornutum*. (A) All elements necessary for the CRISPR-Cas9 genomic editing via bacterial transformation of diatoms are integrated into a single plasmid (pL2-1-Cas9-conjugation-system) using the Golden Gate u-Loop assembly method (Pollack *et al*., 2019). This plasmid is then transferred into *E. coli* bacteria via heat shock and subsequently (B) into *P. tricornutum* via bacterial conjugation. (C) The presence of Cas9 and the two gRNAs on the episome (which persists in the diatom as long as selection pressure is applied) thus enables the generation of mutant lines with an ablation of the targeted lncRNA locus. (D) Successive dilution, purification, and genotyping steps of the transgenic clones then allow the isolation of lines carrying biallelic deletion of the targeted loci. Diagram created in https://BioRender.com.

## Discussion

Long noncoding (lnc)RNAs are transcripts with low to no protein coding potential that have been shown to have important cellular regulatory roles as RNA molecules in eukaryotes [9–11, 13, 15]. Although the great majority of studies characterizing lncRNA function stem from mammalian systems, the functional relevance of several lncRNAs in plants has also been demonstrated. This is the case of *COOLAIR* lncRNAs, antisense transcripts (NATs) that are up-regulated by seasonal winter temperatures and are involved in the mediation of *Arabidopsis thaliana*’s physiological state transition, from non-flowering to flowering, through the coordinated switching of chromatin states of the flowering locus [41]. *IPS1* (INDUCED BY PHOSPHATE STARVATION 1) is a lncRNA that is expressed in response to phosphate depletion and plays a key role in regulating phosphate homeostasis in *Arabidopsis thaliana* [14]. It acts as a molecular sponge, mimicking the target of miR-399 and, due to a mismatched loop at the miRNA target site, instead of being cleaved by the miRNA it sequesters it, reducing its activity towards its actual target (Franco-Zorilla et al., 2007). The discover of *IPS1* and its “target mimicry” action in plants was pioneering in the field of lncRNAs, and was later shown to also occur in mammals [42]. In fact, although lncRNA generally show very little sequence conservation due to lower evolutionary constrains than protein coding genes, there is clear cross-kingdom function conservation [43].

Like metazoans, plants and yeasts, *P. tricornutum* has been previously shown to specifically express lncRNAs in response to environmental cues [4, 19, 21, 23, 24]. Investigating the function of diatom lncRNAs holds great potential to discovering new mechanistic roles of lncRNA, a field that is still in its infancy. A few reports have shown the occurrence and specific expression patterns of lncRNAs in other microalgae, namely in *Chlamydomonas reinhardtii* [17], in *Nannochloropsis oceanica* [44] and in three species of dinoflagellates, *Cladocopium proliferum*, *Durusdinium trenchii* and *Prorocentrum cordatum* [45]. However, taken together, only a handful of reports on microalgae lncRNAs have been published so far. This is not due to the fact that these noncoding transcripts are not present, nor expressed, in marine microalgae but rather that they are still largely overlooked. A recent report on a large-scale screening of transcriptomes originating from 414 species of marine unicellular eukaryotes, from photosynthetic to non-photosynthetic, revealed that all species screened express lncRNAs [20]. In the 101 diatoms screened, lncRNAs represent globally ∼20% of their polyadenylated transcriptomes [20], but their biological relevance remains yet to be resolved. In order to tackle this knowledge gap and enable the investigation of diatom lncRNA functional roles, we sought to develop a method to generate lncRNA knockout (KO) mutant lines, through the use of the CRISPR-Cas9. Since the traditional CRISPR-Cas9 INDEL mutation-induced frameshift in translation and/or premature codon-stop extensively used for the study of protein coding genes will unlikely affect lncRNA function [43], we decided to multiplex our CRISPR-Cas9 approach with a double gRNA strategy in order to induce a large deletion of the targeted locus.

As proof of concept, three *P. tricornutum* lncRNAs responsive to phosphate (Pi) fluctuations to be targeted by our CRISPR-Cas9 approach were selected based on stringent criteria. One of the key criteria we used was that the lncRNA had to be a true intergenic lncRNA. Intergenic lncRNAs originate from genomic regions that are located outside of protein coding gene loci, in both the sense and antisense strands. These can therefore be targeted by the Cas9 endonuclease (which cleaves both DNA strands) and, in theory, avoid the phenotypic misinterpretation caused by the simultaneous editing of a protein coding gene. This is more difficult to achieve with both the intronic lncRNAs and the natural antisense lncRNAs (NATs). Intronic lncRNAs, as their name implies, are lncRNAs that originate from intronic regions of annotated protein coding genes, whereas NATs originate from the opposite strand of protein coding gene loci, overlapping completely or partially with them [21, 43]. Both intronic and NAT lncRNAs are therefore extremely challenging to target by CRISPR-Cas9 since it is almost impossible to not simultaneously affect the cognate protein coding locus and, by consequence, hinder the interpretation of the resulting phenotype.

Another condition used to select *P. tricornutum* lncRNAs to target was that they had to be strong lncRNA candidates and hence have a high probability to be an actual lncRNA. To do that we based our selection on the VotingLNC scores [20]. The *P. tricornutum* lncRNA datasets used in this work [4] were generated before the development of the VotingLNC tool, which is a pipeline that uses 10 diversified protein coding calculator algorithms to define if a transcript is more likely to be coding or noncoding. Potential lncRNAs are considered as such if at least 6 out of the 10 tools assign it as a noncoding transcript, hence have a score of at least 0.6. The three lncRNAs we selected in the present work had scores of 0.8 to 1, which makes them strong lncRNA candidates [20]. It is important to point out that a lncRNA can only be proven as such once it has been shown to be active as an RNA molecule.

Regarding the method to generate functional lncRNA knockouts in *P. tricornutum*, our goal was to develop a plasmid based CRISPR-Cas9 approach where all the necessary parts for the CRISPR-Cas9 multiplexing and transformation via bacterial conjugation, would be combined into a single plasmid. Although the use of multi-plasmid transformation is still widely used for diatom transformation, a growing number of reports have started to focus on developing the use of single plasmids for CRISPR-Cas approaches [28, 31, 32, 46]. The use of a single plasmid avoids the major drawback of multi-plasmid use, namely the increased risk of an incomplete transformation, since it is unlikely that all cells will receive every plasmid, with a significant fraction of transformants lacking components required for a functional CRISPR-Cas9 system. Using the recursive uLoop system, we “domesticated” all the L1 plasmids necessary for the CRISPR-Cas9 double gRNA approach, leading to four different plasmids: pSBL1, the CRISPR-Cas9 and two plasmids containing the sgRNAs of interest. The assembly of these led to a L2 plasmid containing all the necessary parts to perform CRISPR-Cas9 lncRNA locus ablation in *P. tricornutum* by bacterial conjugation (Figure 1). The advantage of this assembly resides in the ease to generate an effective plasmid for a selected gene. Since all the components now exist at the L1 level, a simple PCR is sufficient to replace the guide RNA sequences (as described in Material and Methods) hence, with two PCRs (one for each of the L1 gRNA plasmids) and one uLoop assembly, a ready to use a new L2 plasmid to perform a multiplexed CRISPR-Cas9 strategy by bacterial conjugation in *P. tricornutum* can be easily obtained to target any locus of choice.

The present protocol has been developed with the goal to perform functional studies on lncRNAs in *P. tricornutum*, however, it can also be easily adapted to other diatoms, or other microalgae, by simply replacing the *P. tricornutum FcpB* promoters by species-specific strong endogenous promoters. The main bottleneck to generate lncRNA knock-out mutants in other diatoms, or other microalgae, will be the transformation step. Although the number of diatoms with transformation protocols available has been increasing in recent years [47], *P. tricornutum* is still the diatom with the largest number of transformation reports. Different approaches are currently available to transform *P. tricornutum* namely biolistic bombardment [48], bacterial conjugation [39], electroporation [49–51] and, more recently, through the use of polyethylene glycol (PEG) after the enzymatic digestion of the cell wall [51]. The first two methods are still the most commonly used transformation approaches used with *P. tricornutum* and, in this work, bacterial conjugation was chosen to generate lncRNA KO mutant lines over biolistic for several reasons. Bacterial conjugation is a very simple method to use, requiring no special expensive instruments, nor elaborate protocols, and has the highest transformation efficiency rate (4 transformants per 10^4^ cells, which is up to 1000 times higher than biolistic) [39, 52]. Also, episome DNA delivery to diatoms avoids the side-effects generated by the biolistic approach such as the random, multi-copy integration of the recombinant DNA in the genome and, of significance importance, it also enables the curation of the episome by removal of the selection pressure [39, 46, 52, 53]. Curing of the episome enables avoiding critical issues related to excess Cas9 concentration in the cell, namely the increased occurrence of off-target effects [54]. It also allows to obtain marker free mutated diatoms which, in our case eliminates the need to use zeocin in the culture media to maintain the mutant strains.

After transformation, the first colony-based PCR screening of the exconjugants revealed that 11% of the colonies screened presented a biallelic full ablation of the lncRNA loci (Fig. 3). This shows that in these colonies, both multiplexed gRNAs had successfully simultaneously cut the targeted regions in both homologue chromosomes. Similar results were reported in another multiplexed study using two gRNAs to target two different protein coding genes [28]. Since our goal was to obtain biallelic mutants harboring a full ablation of the targeted lncRNA loci, we did not investigate further the single editing events that had occurred. Also, we cannot exclude the occurrence of off-target effects in the isolated biallelic mutants, although this has been shown to not have a significant occurrence in *P. tricornutum* when using bacterial conjugation and episome curing [55].

Preliminary experiments assessing specific growth rates under phosphate fluctuation revealed that one of the lncRNA ablated mutants, *Δlinc24*, showed a growth phenotype, dividing significantly faster under full media and under Pi stress than the WT (Fig. 4). This lncRNA has been characterized as being up-regulated by phosphate depletion but, interestingly it is also up-regulated under nitrogen depletion [4]. This suggests that rather than being related to a specific nutrient stress, namely Pi, *Linc24* is likely involved in a general nutrient depletion response. A common response to both Pi and N depletion in *P. tricornutum* is growth arrest [4, 56, 57]. The fact that *Δlinc24* shows a growth phenotype, namely increased growth rates when compared to the WT (Fig. 4), suggests that it could be involved in the regulation of cell division under nutrient stress. More evidence is now needed to confirm this, namely repeating the growth experiments with different nutrient fluctuation conditions. Furthermore, it would be important to validate the findings with another genetically independent mutant line, harboring the *Linc24* locus ablation. The two other lncRNAs mutants generated, *Δlinc77* and *Δlinc91*, did not show a growth phenotype under the conditions tested (Fig. 4). This can either mean that these two loci have no biological relevance or of the existence of regulatory redundant pathways that enable *P. tricornutum* to compensate its growth under Pi fluctuations, despite the impact of the lncRNA ablations. In order to thoroughly assess the existence or the absence of the biological relevance of *Linc77* and *Linc91* in *P. tricornutum*, other physiological phenotyping is required, namely evaluating their photosynthetic efficiency and capacity under nutrient fluctuations.

When assessing lncRNA function by reverse genetics, it is important to note that the ablation of a lncRNA locus will not elucidate if the resulting phenotype is due to the absence of the transcript or of the act of transcription itself. The existence of a phenotype in the lncRNA ablated mutants will nonetheless reveal the importance of the noncoding locus as a functional genomic element [33]. Other, orthogonal experiments are needed to provide insights into the existence of mechanistic roles, both *in vitro* and *in vivo*. Namely, doing simple rescue experiments by complementing the mutant line with the lncRNA WT locus and study the resulting phenotype could provide hints into whether the transcript is active as and RNA molecule acting in *trans*.

## Conclusions

In conclusion, we successfully generated a simple and efficient protocol which enabled the generation of the first CRISPR-Cas9 ablated lncRNA mutants in the diatom *P. tricornutum*. To our knowledge, this work represents the first report of the generation of lncRNA knockout mutants in a microalga. We believe that this protocol will be extremely useful to perform lncRNA functional studies in *P. tricornutum* and beyond, enabling to shed the first insights on the involvement and the roles of lncRNAs in diatom resilience to a fluctuating environment.

## Acknowledgments

The authors wish to thank Christopher Dupont for kindly providing the genetic parts from the uLoop library as well as Vincent A. Bielensky and Bernardo Pollack for the discussions regarding uLoop domestication.

## Authors’ contributions

HCC conceived and designed the research; FC and NR performed the molecular experiments; HM performed the mutant growth experiments; HCC wrote the manuscript with inputs from FC, NR and HM. All authors read and approved the final manuscript.

## Availability of data and materials

The plasmids generated on this study are available upon request to the corresponding author. The detailed step-by-step protocol generated in this work is available at protocols.io under https://dx.doi.org/10.17504/protocols.io.kxygx3yyog8j/v1.

## Supplementary data

Supplementary Figure S1 - PCR screening of the exconjugants

Supplemental Table S1 - Primer sequence information

Supplementary File S1 - Sanger sequencing of the CRISPR-Cas9 KO mutants

