## Supplementary figures and images for "A quick and efficient method enabling lncRNA functional studies in the marine diatom *Phaeodactylum tricornutum* using a multiplexed CRISPR-Cas9 editing approach"

### Supplementary Figure S1

Supplementary Figure S1: PCR screening of the exconjugants.

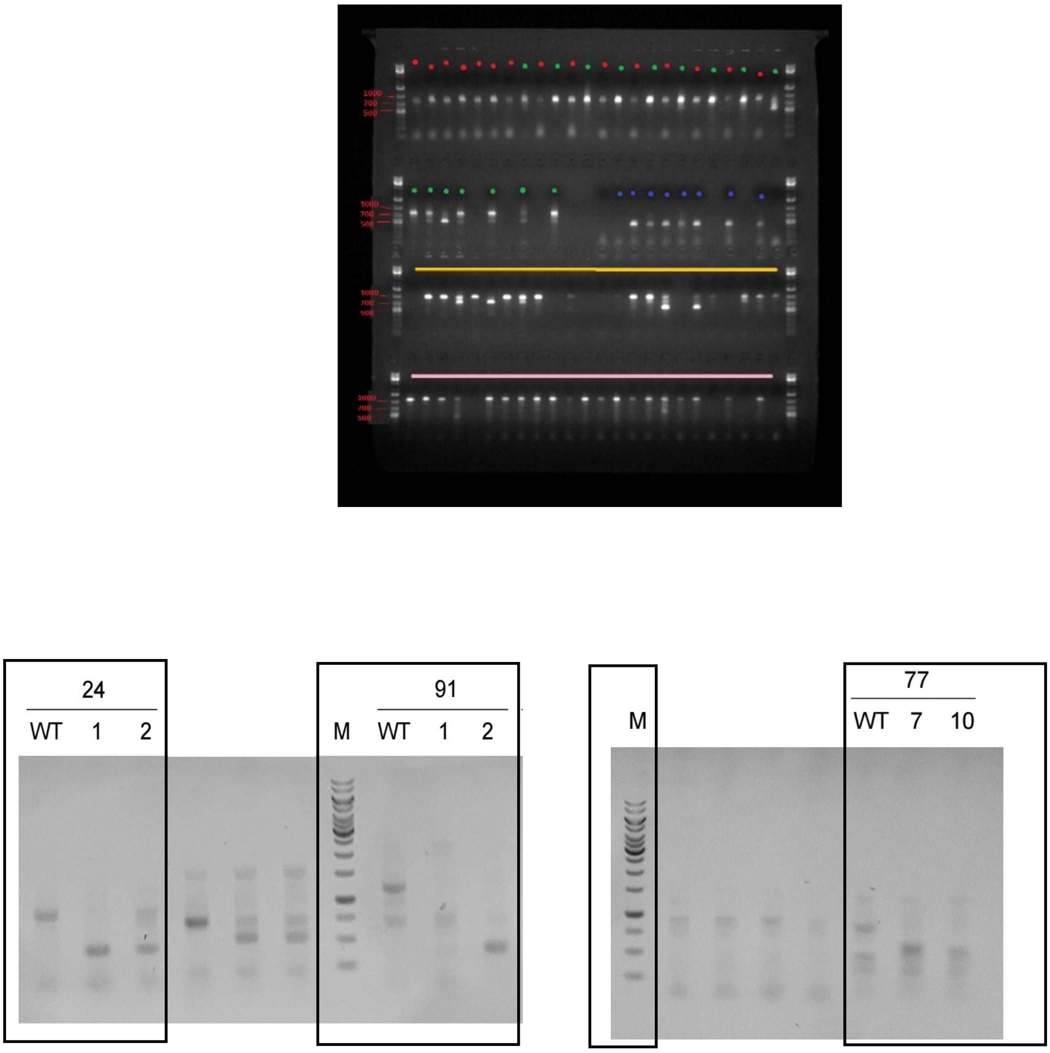
