## Supplementary File S1 for "A quick and efficient method enabling lncRNA functional studies in the marine diatom *Phaeodactylum tricornutum* using a multiplexed CRISPR-Cas9 editing approach"

### Mutant Sanger sequencing data

*ΔLnc24*: Δ436 nt (total 502 nt), ablation ~87% (86,9%)

GCGCGACTACCTAAGTC**AG**ctcctcgccggatttcgtgagtacaagattgtctcacgggatcgatcaatcccattcctgcccctctcccacga
cacgaaatcgtcctttttatggttaggtatcgagaagagcttggttgcacacggaatttcatgtgtggaatacgatcgaccgatagtgatacctat
acattcccaaagggttctactctcttgcgaagagtggggggtgaggaacggtattggcaccggacggagtcgattttgacggggagtcctccgta
aaccttacgttgcccggcacctctcggctctacttttacgaattgatgaaaacacgtctcgatgcgcagattgatcttcaggcgagtgtcactgtga
gagcgtggtattcactttactagagagtctgctagagagagagaaatgcccgacccaatttcacataagcagt**AA**CTATAGATAGGCGAC
TGTTTTCGCGCAAAAGTCGTGACGCGTCCC

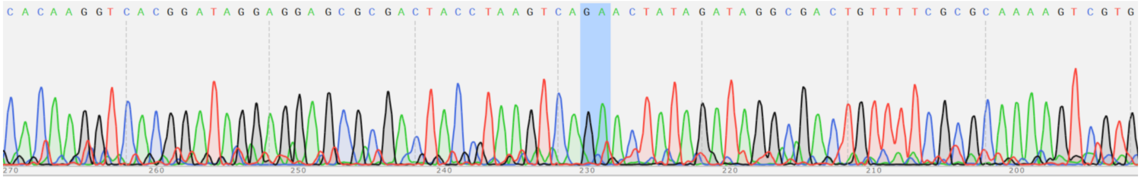

*ΔLnc77*: Δ268 nt (total 454 nt), ablation ~60% (59%)

GCATCGGCGACTCGACAACGACACGAAGAGCCAAGTATTTCTCCTCTACCTCCCCAGGCTCTATCAGAGCTGGCAAT
CTCTTCAAACGCTCGAGCACAACGAAAACGCTACCTTTGATCCC**GC**tactaggaaaatgcgacaataactggctacaatcgtgc
ctaagagaccactggtctgccgtgatcaaaggcgtaaaccataaaagttatagacttgtccatgtcagggaaccactgttcggtggttgacttcca
ccgccaatatctccttacggtcatcgtccccacacaatgaccggccgtcatcaaactagcgtattcaactaaccttacaattttaggatctacagaa
accaaaaggtctgtttctctaggatagttggtcct**ga**TACATTCTATCGTCTAATTAAGACCAACTCACTCCCTTGAGGAGCTCTC
CCCTTTCTCAACGC

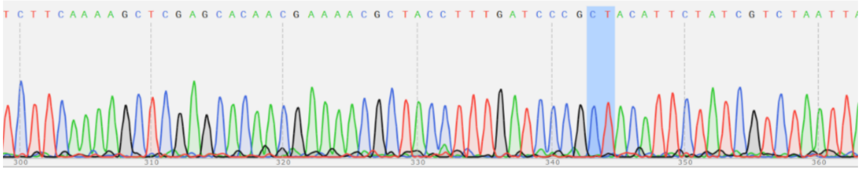

*ΔLnc91* - Δ812 nt (total 906 nt), ablation 90% (89,6%)

CAAATTTCAAGACTCCCACAAGCCACCT**ACG**tattatgccttacagatagaggacataggtttctattatccttccagagaggcattcat
caggtctcgcaccgcaagggaggattgactgtccaaggaatgcctgctaaagaatcctgtacaattgtgtggaggaatcaaactctagctttcga
aacgatttagcagtacactcactgacagtaagtgacttgtagcctaatagctcagctcaagtatcgtcgaaacacgctccaatgaaatttattata
atacattataaaaaacagacttccttcctacactgattgcgcaccggaattctccgagtgaggtatagagaatgttaatgtgggaaaacaggaac
agccacggtatcaagagagtttgtgttgtttgcccaagtgacaaaaggcagtaaagatatatgggcttgacgattccggccagaccaaagcatg
ggagcaataggtattgcaagtcaaattactgggaatacgatcttcccgcggtgtcgctaactgtgagtgaattgatggaaataaatgagcaaac
aaaaaaacagaacgaatctttttagcgtttgatgcagtgccaccatactcggggctatggatattggtagacggcaagaattgacaagaagttt
gtttcagttattttgtttttcgttgcggttagacaatcacttcttcgtatgttaacagtcaactttgttactccgtcttcactgcttgtttccggtgtcttcgt
gtaattcggtcaaatcaaatatactccgtaggaaaaaatcagtcgtttctgaaaatgacgaggaacgttccctatttatag**ATG**TCTTACGAC
TATAGTATTTAAATCAAAACGAATAATTGCAGTCGGACGCTTCTGGCTGTC

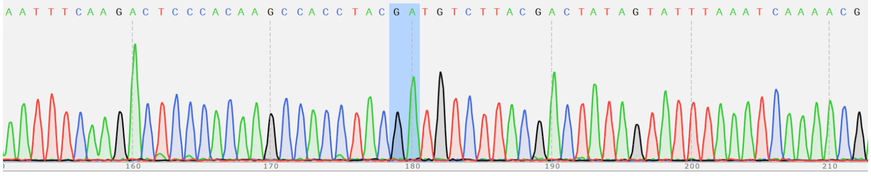

Red: PAM | Yellow highlight gRNA | Bold: where **Cas9 cut** | Lower case: what is absent in the biallelic mutants
