## Supplementary Table S1 for "A quick and efficient method enabling lncRNA functional studies in the marine diatom *Phaeodactylum tricornutum* using a multiplexed CRISPR-Cas9 editing approach"

Supplementary Table S1 - Primer sequence information for PCR screening of exconjugar

| LncRNA | Primer | Sequence |
|---|---|---|
| ***Linc24*** | ΔFwd | CGTGCTGGTCTGGAAGGAAT |
| | ΔRev | CGATGGGACTGGCAGTGAAT |
| ***Linc77*** | ΔFwd | CCTGACGATAGAGGCTCGAC |
| | ΔRev | TCTGTATCGTTTCGCGCTCA |
| ***Linc91*** | ΔFwd | ATGTGCAGCCAGGACTAACC |
| | ΔRev | ATTTCGTGGCACTTTCGCAG |

ıts
